# Time-series Classification for Patients under Active Surveillance and Screening Using Echo State Network

**DOI:** 10.1101/2023.04.24.538057

**Authors:** Zonglun Li, Alexey Zaikin, Oleg Blyuss

## Abstract

Over the past few decades, more and more patients come on follow-up studies such as active surveillance and screening, which results in a vast amount of time-series data in the health department. Each Patient typically has a small but different number of visits to the doctor and the time interval between the visits is heterogeneous. Nowadays, many machine learning tasks in relation to time series data are carried out using deep recurrent neural networks (RNN). However, deep neural networks consume enormous computational power as all weights in the network need to be trained through back-propagation. Conversely, echo state network (ESN), another form of RNN, demonstrates low training cost and the potential of it is still largely untapped. Therefore, in this article we will develop a new methodology that can classify aforementioned time-series data using the echo state network. We will also discuss how to address the heterogeneity in the time interval arising from the data of this type and how our model can also potentially fit other time-series data.

## 1 Introduction

Data explosion gives rise to data proliferation of various types and formats. Among them, time-series data have played an increasingly vital role in medical research such as cancer and diabetes, not least with the rapid development of machine learning techniques Kiriu et al. [2018], Lam et al. [2018], Whitwell et al. [2020], Gentry-Maharaj et al. [2020], Sushentsev et al. [2023], Alhassan et al. [2018], Xie and Wang [2020]. Nowadays, various clinical programs have been deployed worldwide to either facilitate the early diagnosis, or monitor the progression of the disease by inviting the population at risk to get tested on a regular basis. Among them, active surveillance and screening are part and parcel to the wellbeing of patients and sustainability of the health system for Disease Control et al. [2012], Duffy et al. [2020], Blyuss et al. [2023], Cooperberg et al. [2011], Klotz et al. [2015], Sushentsev et al. [2022]. Active surveillance is a treatment plan for low-risk cancers that involves monitoring the status of a patient without providing any treatment until it becomes necessary (e.g., when the progression occurs). This can substantially reduce the side effect on patients as well as the cost on the medical resources. Cancer screening is a test that aims to seek early sign of cancer before the symptom appears when the medical intervention is more likely to be effective. It has now become a nationwide program in many western countries. In spite of the disparate natures of these techniques, they share a number of similarities in terms of the data structure. Patients typically have multiple visits to the clinicians and have their samples taken that generally spans years. It therefore results in short time-series data with the expression of bio-markers recorded at every time point. Furthermore, the diverse behaviors of individual patients cause them to have different times of visit and the visits are not evenly spaced.

Lately, deep recurrent neural networks such as long short-term memory (LSTM) have been extensively employed for cancer prediction and detection Dutta et al. [2020], Budak et al. [2019], Begum et al. [2022], Sushentsev et al. [2023]. However, although they can largely yield impressive performance scores such as accuracy and ROC AUC, the training cost is exceptionally high down to the fact that the training requires the back-propagation from the last layer to the first layer where all weights in the network need to be adapted. Reservoir computing is a type of recurrent neural network (RNN) that maps input signal to a non-linear high-dimensional dynamical system Jaeger and Haas [2004], Tanaka et al. [2019], Schrauwen et al. [2007], Nakajima and Fischer [2021]. Since the neurons in the reservoir layer are connected recurrently, it can in theory, memorize relatively long short-term memory. As opposed to the artificial neural network, only the output layer in a reservoir computer needs to be trained and all the other connections are fixed and this gives reservoir computing the edge of high efficiency as compared to its counterparts. Over the last two decades, echo state network, a class of reservoir computing where the state of neurons in the reservoir layer is updated by a non-linear activation function, has been applied to time-series learning in various fields Lin et al. [2009], Li et al. [2012], Song et al. [2020], Li et al. [2020], Liu et al. [2022], Shahi et al. [2022]. Nevertheless, most of them focused on the prediction of time-series and the development of classification methodologies is still lacking. Although several studies have already touched on this topic Skowronski and Harris [2007], Ma et al. [2016], Wang et al. [2016], Stefenon et al. [2022], the research along this line has multiple concerns when it comes to the application on the aforementioned data. Most importantly, all of these approaches are either generic or are designed for some particular applications and do not deal with the unique feature of our data. Hence, in this article, we will propose an intuitive methodology that can properly accommodate the unique features of the data and exhibit its potential to become a generic approach.

## 2 Datasets

In this study, we assess the performance of our methodology using two datasets and we call them radiomics data and BD data respectively. The radiomics dataset (Sushentsev et al. [2023]) contains 76 patients on active surveillance for prostate cancer. The predictors (features) of interest are PSA density and 44 radiomics features. The BD dataset (Blyuss et al. [2015], Marino et al. [2017], Vázquez et al. [2018]) contains 222 patients on screening for ovarian cancer after removing those with only one visit. The predictors are CA125, Glycodelin, HE4, MSLN25, MMP745 and CYFRA55 respectively, among which CA125 is regarded as the primary predictor.

## 3 Methods

Suppose *k* features (markers) of *N* patients have been taken for analysis and patient *i* has *n*_*i*_ time points (number of visits). We denote the input vector of patient *i* at time point 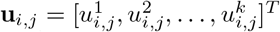, where *i* = 1, …, *N* and *j* = 1, …, *n*_*i*_. The outcome of patient *i* is denoted by *y*_*i*_ which is either 0 or 1.

Figure 1 displays the echo state network that we used for our study. The dynamics is governed by Equation 1. **x**_*i,j*_ represents the internal states for patient *i* at time point *j*, **W**_*in*_ represents the input weight matrix and **W**_*res*_ represents the weight matrix in the reservoir layer, and *a* denotes the leakage rate. Unlike the artificial neural network, here both **W**_*in*_ and **W**_*res*_ are fixed throughout the training.

**Figure 1.**
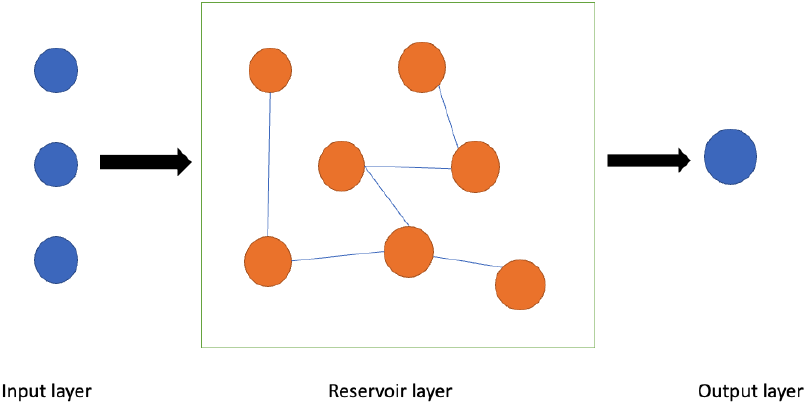
Echo state network.

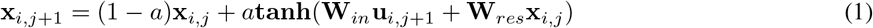

Assume that there are *k* neurons in the input layer, *M* neurons in the reservoir layer and one neuron (for classification) in the output layer. For each input vector **u**_*i,j*_, we pass it from the input layer to the reservoir layer and collect all internal states in the reservoir layer 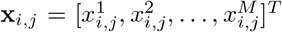 and we will explain in due course how to train the output layer.

In general, the datasets of concern do not have sufficient samples to carry out a more unbiased evaluation by splitting them into training and test sets, partly because the patients participate on a voluntary basis and the project requires consistent follow-up over several years. Therefore, here we implement k-fold cross-validation on the entire dataset to evaluate the performance of our method.

For the BD data, we use 5-fold cross validation on the entire dataset in order to evaluate the performance with *N*_*train*_ denoting the size of the training set and *N*_*test*_ the size of the test set in each iteration, *N*_*train*_ + *N*_*test*_ = *N*. In the training stage, for each patient *i*, we stack all internal states according to the time sequence. Note that the internal states will need to be reset to zero before processing a new patient. Then we stack the aforementioned internal states of all patients and the corresponding outcomes. The resulting stacked matrix for internal states will be

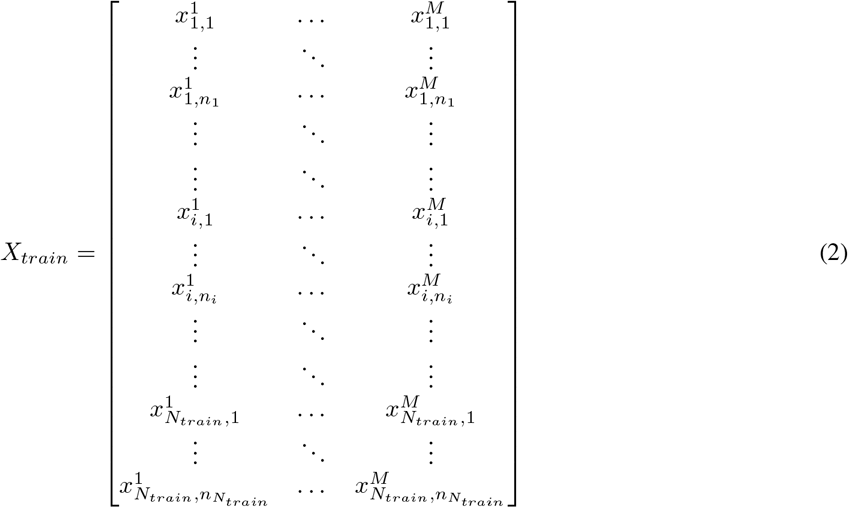

and the corresponding stacked output vector will be 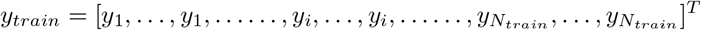. In a nutshell, if patient *i* has *n*_*i*_ time points, then there will be *n*_*i*_ entries in *y*_*train*_ and *n*_*i*_ rows in *X*_*train*_. One can think of *X*_*train*_ as the design matrix in the regression model. This is to ensure that we have enough samples to train the classifier while taking into account the temporal dynamics. By training the internal states that incorporate diverse lengths of history (not just the last one) of a particular patient, it also potentially enables a better generalization when seeing the longitudinal features from other patients. Then we fit *y*_*train*_ to *X*_*train*_ using the linear support vector machine (SVM) to train the classifier (output layer) as it is effective in high dimensional space, not least when the number of samples is not sufficient enough. In the testing stage, we collect all internal states of the test set in a way identical to Equation 2, only with *N*_*train*_ being replaced by *N*_*test*_. Then we employ the trained classifier to predict the outcome (in probability) of 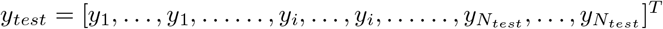. At the moment, each patient has multiple predicted outcomes as explained previously. In order to compute the performance score (e.g., ROC AUC), one needs to convert them into a single outcome. One option is to select the prediction at the last time point to be the representative of the particular patient. In this way, we put more emphasis on the last time point which will potentially give rise to a better score, even though the internal states of the last time point is dependent on the previous time steps and the classifier has been trained on all time points. Finally, we compute the average of the scores obtained in each iteration and utilize it to reflect the goodness of our model.

For the radiomics data, we use leave-one-out cross validation (LOOCV) to evaluate the performance due to the sparsity of samples. The procedure is similar to that of the BD data except that now each iteration only has one sample (patient) in the test set. Instead of computing the score of the test set, we store the representative outcome (e.g., the last point) of that particular patient in each iteration. Hence, we will be able to collect the predicted outcomes of all patients and compute the performance scores accordingly.

## 4 Extension: How to include sample taken time?

Thus far, we have ignored the sample taken time for each patient. However, take for instance a given patient has samples taken at month 1, 11 and 12, then it may not be plausible to assume that the features at these points can be directly used as input time-series data to the reservoir computer as it cannot differentiate how apart these time points are. Unfortunately, traditional RNN is not able to capture the difference in time interval automatically. Zhu et al. [2017] and Gao et al. [2019] proposed two variants of LSTM to deal with heterogeneous time intervals. However, Zhu et al. [2017] features the recommendation systems and the essence of the time interval is fundamentally different from the one in our context; Gao et al. [2019] adds a temporal emphasis function to the forget and the input gate but does not handle the time interval explicitly. Not to mention that the architecture of LSTM cannot be translated to ESN. Therefore, in this section we will develop a methodology that can properly handle the heterogeneity in the time interval in the backdrop of active surveillance and screening.

One thing we need to bear in mind is that the disease progression is largely a continuous process and one may consider introducing artificial data to fill the gap of wide intervals. Here we introduce a scaling factor ∆*t* such that given the actual sample taken times (after being converted from dates to real numbers) 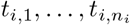 for patient *i*, we further transform them into integer numbers with

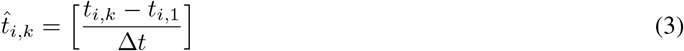

for *k* = 1, …, *n*_*i*_, where the bracket symbol rounds the fraction to the nearest integer. In other words, if two consecutive time points are equal, 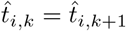, then one of the data at this time point will be removed; if 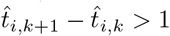, then we need to add artificial data in this interval. For instance, after the transform, if 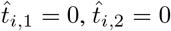 and 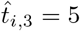, then the feature record at *t*_*i*,2_ will be removed and we will need to introduce four more artificial feature records at time point 1,2,3,4. One way to generate these artificial data is to calculate the difference of feature expressions at two consecutive recorded times (0 and 5 in this case) and spread it over the interval. For example, if the original time-series data for a patient with only one marker is

|  |  |  |  |  |
| --- | --- | --- | --- | --- |
| time | 66 | 66.7 | 68.2 | 72 |
| marker expression | 10 | 10 | 15 | 25 |

where the first row is the sample taken time and the second row is the corresponding marker expression. After transforming with ∆*t* = 2, it turns out to be

|  |  |  |  |  |
| --- | --- | --- | --- | --- |
| time | 0 | 0 | 1 | 3 |
| marker expression | 10 | 10 | 15 | 25 |

Then the input sequence to the reservoir computer will be 10, 15, 20, 25 as we remove one value at time 0 and add an artificial value at time 2.

Besides making the time-series data more homogeneous, it can also generate more complex behaviors if we choose a small ∆*t*, because in this way we prolong the length of the internal states for each patient. However, it will therefore increase the training cost since more samples need to be trained and we introduce another parameter ∆*t* that needs to be explored in order to select the best model.

## 5 Results

In this section, we will report the best score among a variety of parameter combinations in each study case. Without the inclusion of the sample taken time, *M* and *a* will be searched to report the best performance; with the inclusion, *M, a* and ∆*t* will be explored.

### 5.1 Without including sample taken time

For the BD data, we evaluate the performance of the following two cases and the results are shown in Table 1.

**Table 1:** BD data

| Model | ROC AUC |
| --- | --- |
| 1 | 0.952 |
| 2 | 0.963 |

1. CA125 only.
2. CA125, Glycodelin, HE4, MSLN25, MMP745, CYFRA55.

For the radiomics data, we evaluate the performance of the following three cases and the results are shown in Table 2.

**Table 2:**
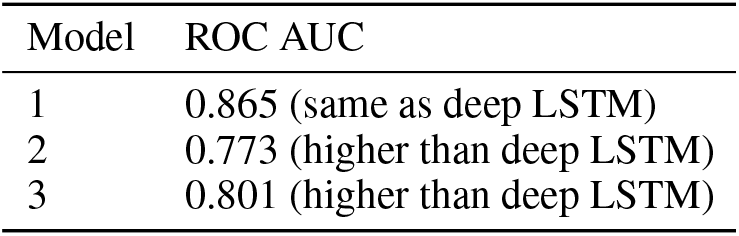
Radiomics data

| Model | ROC AUC |
| --- | --- |
| 1 | 0.865 (same as deep LSTM) |
| 2 | 0.773 (higher than deep LSTM) |
| 3 | 0.801 (higher than deep LSTM) |

1. PSA density and all radiomics features.
2. PSA density only.
3. All radiomics features on

### 5.2 Including sample taken time

For the BD data, we evaluate the performance of the following two cases and the results are shown in Table 3.

**Table 3:** BD data

| Model | ROC AUC |
| --- | --- |
| 1 | 0.951 (same as not including) |
| 2 | 0.974 (slightly higher than not including) |

1. 1, CA125 only.
2. CA125, Glycodelin, HE4, MSLN25, MMP745, CYFRA55.

For the radiomics data, we evaluate the performance of the following two cases and the results are shown in Table 4.

**Table 4:** Radiomics data

| Model | ROC AUC |
| --- | --- |
| 1 | 0.800 (higher than not including) |
| 2 | 0.800 (same as not including) |

1. PSA density only.
2. All radiomics features only.

Here we do not consider the case of PSA density and all radiomics features because the sample taken times of them are different.

## 6 Discussion and Future Work

In this work, we developed a methodology that can predict the outcomes of patients on active surveillance and screening using ESN with high efficiency and desirable performance. The methodology can also be easily generalized to other time-series data with a similar structure. As previously mentioned, data collected from active surveillance and screening typically have small sample size and short time-series. Nevertheless, the method is adaptable when it comes to other longitudinal data with much longer time-series. Instead of placing all internal states into the design matrix 2, one can sample them with skip in order to reduce the training cost. For instance, if the corresponding internal states for patient *i* are **x**_*i,j*_, *j* = 1, …, *n*_*i*_, then one can choose 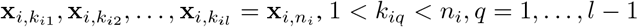 to form the design matrix. Even if we will miss some internal states at some particular time points, it can still capture the dependency among different time points since **x**_*i,j*_ will keep track of the history before time step *j*. Hence, it looks promising to develop a more generic approach for time-series data classification and assess the model using a wider spectrum of data.

Besides, more thoughts can be given as to how to improve the model and make it in more compliance with the existing theories. Thus far, the first input vector of patient *i*, **u**_*i*,1_ has been used as the first input to the reservoir computer. Some previous studies have shown that the ability of the dynamical system to achieve the desired outcome is optimized near the edge of chaos Legenstein and Maass [2007a,b], Boedecker et al. [2012]. What if we introduce a burn-out period before the longitudinal features for each patient which can potentially drive the dynamics in the reservoir layer near this critical state when seeing the features of interest? Additionally, so as to address the challenge of the heterogeneous time interval, we adopted a heuristic approach to ensure that the inputs to the reservoir computer are more evenly spaced. Lately, several neural ODEs models have been developed to account for irregular time series Rubanova et al. [2019], Kidger et al. [2020], Schirmer et al. [2022]. It might be interesting to see if we could integrate the ideas along this direction into our current framework and deal with the irregularity more properly.

## 7 Acknowledgement

We acknowledge support from CRUK grant EDDCPJT/100022 and CRUK ACED grant EOCEDAAP/100009.

